# Species matter for predicting the functioning of evolving microbial communities

**DOI:** 10.1101/666685

**Authors:** Timothy Giles Barraclough

## Abstract

Humans depend on microbial communities for numerous ecosystem services such as global nutrient cycles, plant growth and their digestive health. Yet predicting dynamics and functioning of these complex systems is hard, making interventions to enhance functioning harder still. One simplifying approach is to assume that functioning can be predicted from the set of enzymes present in a community. Alternatively, ecological and evolutionary dynamics of species, which depend on how enzymes are packaged among species, might be vital for predicting community functioning. I investigate these alternatives by extending classical chemostat models of bacterial growth to multiple species that evolve in their use of chemical resources. Ecological interactions emerge from patterns of resource use, which change as species evolve in their allocation of metabolic enzymes. Measures of community functioning derive in turn from metabolite concentrations and bacterial density. Although the model shows considerable functional redundancy, species packaging does matter by introducing constraints on whether enzyme levels can reach optimum levels for the whole system. Evolution can either promote or reduce functioning compared to purely ecological models, depending on the shape of trade-offs in resource use. The model provides baseline theory for interpreting emerging data on evolution and functioning in real bacterial communities.

## INTRODUCTION

Many biological processes that humans depend upon – such as global nutrient cycling, plant growth and digestive health – rely on the action of microbial communities with hundreds or even thousands of species [1–4]. A key challenge is therefore to understand such complex systems in sufficient detail to be able to predict overall functioning and how it changes under fluctuating conditions [5, 6].

In both microbes and eukaryotes, measures of functioning such as yield or productivity tend to increase towards an asymptote as species diversity increases [7]. However, the mechanisms for these effects, in terms of resource use of species, interactions among species and their dynamics over time, remain largely unknown [8]. Models are needed to identify key features that determine community functioning, and how we can modify them to enhance function [9, 10].

The bacteria in our digestive tracts are a notable case [11]. Outnumbering human cells and containing 300 times as many metabolic genes as the human genome [12], the gut community breaks down indigestible macromolecules in our diet and releases beneficial compounds [3, 13]. The resulting metabolites provide a source of energy, especially in non-industrialised diets, and feed into numerous physiological pathways, including appetite regulation and immune responses [14, 15]. Disruption of the gut microbiota and of metabolite production leads to gut disease and impaired health.

Despite the wealth of new data coming from metagenome sequencing, it remains unclear how beneficial functions of microbiomes depend on the action of constituent species. Some authors argue for widespread functional redundancy and that a far smaller set of species could perform adequate function [16, 17]. Alternatively, full diversity might be vital for overall functioning, if species have specialized metabolic roles and provide robustness of functioning in varying conditions such as fluctuation in diet [18]. Metabolic function of component species can be inferred from genome sequences [19] or from culture or co-culture experiments [20]. The hard part is to map from species metabolism and growth to overall community functioning.

One simplifying approach is to infer function from metagenomes without considering how genes are packaged into species [12]. Many animal and plant ecologists would be skeptical of predicting ecosystem functioning from a homogenized blend of species traits. Although the metagenome provides a snapshot of the metabolic diversity [21], population dynamics and hence relative abundances of different functions depend on whole genomes and their growth in a community of species. However, flux balance approaches for modeling metabolism of single cells are useful without considering internal metabolite and enzyme dynamics in detail, and recently these have been applied to communities [22, 23]. Perhaps species densities shift to produce concentrations of enzymes at steady-state that are the same at the whole-system level irrespective of how those enzymes are packaged among species.

Another issue for predicting functioning of microbial communities is evolution of component species [24]. In productive communities such as the gut, species are likely to evolve over short timescales (weeks) in response to changes in physical environment, resource availability, or species interactions [25, 26]. Evolution in resource use might alter overall functioning of the community. For example, if species evolve to partition resources or to increase their rate of use of metabolites, this could enhance degradation of input resources [27]. Alternatively, evolution might reduce functioning if species evolve to degrade intermediates that are beneficial for overall functioning. Natural selection operates on individuals rather than on the whole community, so evolution does not necessarily optimize ecosystem functioning. The interplay between ecological and evolutionary responses should therefore determine whether functioning can be predicted from a metagenome type approach or whether a species-based approach is required.

I address these questions using a model of species metabolizing input resources via a series of intermediates (Fig. 1). The model is an extension of standard chemostat models of bacterial population growth [28–30], adding evolution of resource use of each species. It is designed to use parameters that could be measured in experiments with real microbes, but simplified to enable general understanding of such systems. Although many additional processes operate in real systems, the model provides baseline predictions for evolution and functioning in real bacterial communities. After illustrating how the model can be used to predict different aspects of functioning, I evaluate the effects of packaging the same enzymes in different ways among species (focusing on the comparison of specialists and generalists), in both non-evolving and evolving versions.

**Figure 1.**
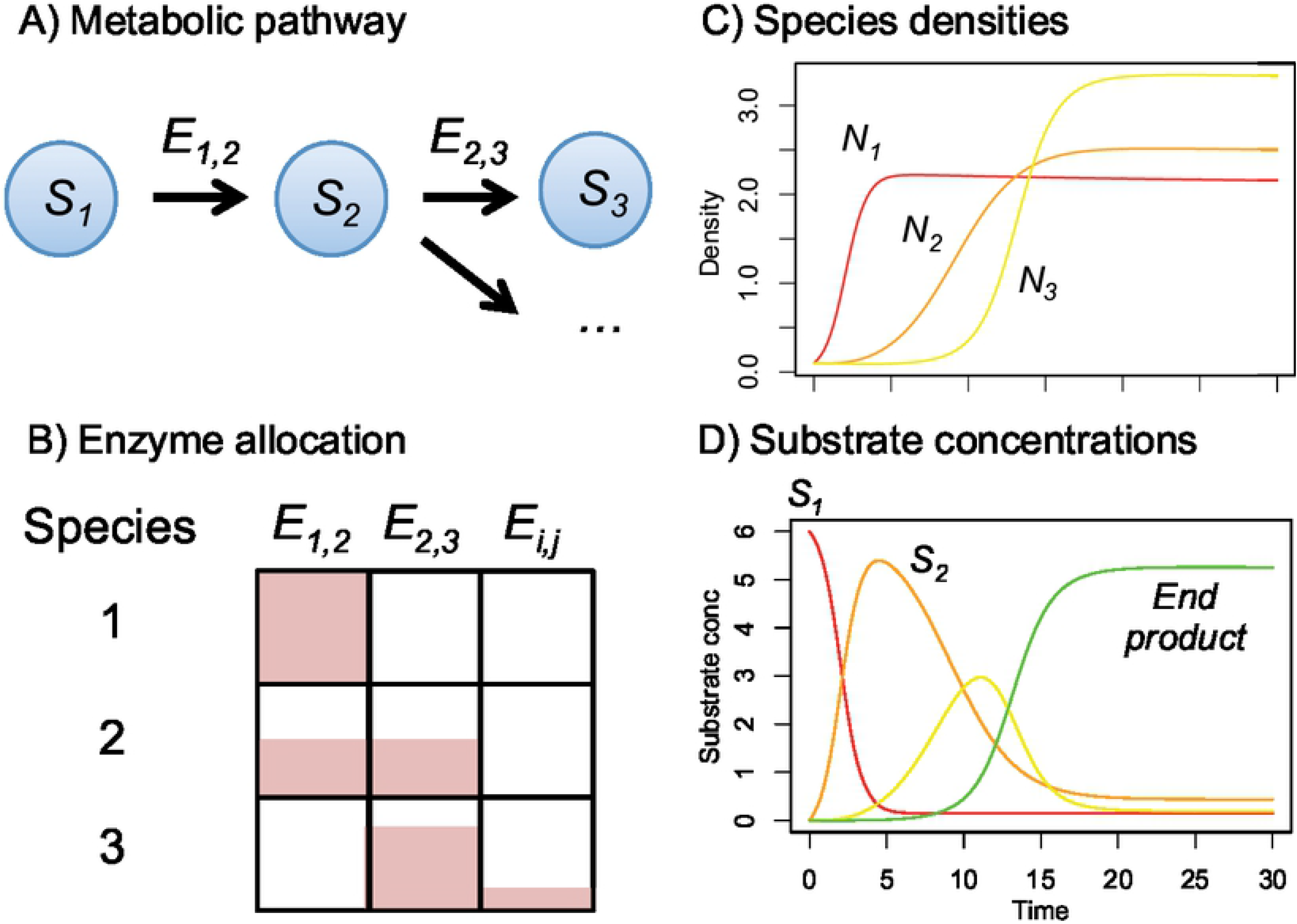
Schematic diagram showing components and outputs of the model. Multiple coexisting species vary in their ability to metabolise substrates in a metabolic pathway. A) Substrates are connected through a pathway performed by a set of enzymes: E_ij_ metabolises i to j. B) Resource use of each species is specified by a matrix of enzyme allocations: species can either be specialist on one substrate or generalists on multiple substrates. C) Species grow in proportion to the amount of substrate they metabolise and D) substrate concentrations change as they are degraded or produced by one or more species. From these variables, various aspects of community functioning can be calculated, such as the rate of degradation of input resources. In the evolutionary model, the enzyme allocations also evolve over time.

## METHODS

### Ecological model with no evolution

Consider *n* species with densities *N*_*k*_ *(k*=*1,2,3,…,n)* growing in an unstructured environment via metabolism of *m* substrates with concentration *S*_*i*_ (*i*=1 to *m*). There is flow through the system at a constant dilution rate *D* (chemostat conditions). Some substrates are present in the inflow (at concentration *Q*_*i*_) whereas others are absent and generated by metabolism of inflow substrates. Both substrates and bacteria are removed in the outflow.

Growth occurs by metabolizing substrates according to a Monod growth function [29]. The rate per cell of converting substrate *i* into substrate *j* in species *k* is

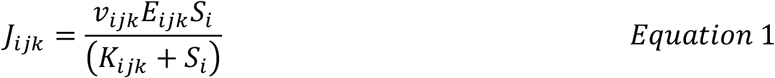

where *v* is the maximum reaction rate per molecule of enzyme (sometimes called *k*_*cat*_), *E* is the number of molecules of enzyme per cell, and *K* is the substrate concentration at which reaction rate is half its maximum value (i.e. affinity for the substrate is 1/*K*, all parameters defined in table 1). In order to explore additional analytical solutions, I also considered a simpler model with a linear growth function, i.e. replacing *J*_*ijk*_ = *v*_*ijk*_*E*_*ijk*_*S*_*i*_.

**Table 1.** List of state variables and parameters

| Term | Definition | Hypothetical units |
| --- | --- | --- |
| $N_k$ | Density of species $k$ | Cells ml<sup>-1</sup> |
| $S_i$ | Concentration of substrate $i$ | mmol ml<sup>-1</sup> |
| $E_{ijk}$ | Allocation of species $k$ to enzyme metabolising substrate $i$ to substrate $j$ | Molecules cell<sup>-1</sup> (relative to total enzyme content, assumed fixed) |
| $v_{ijk}$ | Maximum reaction rate | mmol molecule<sup>-1</sup> enzyme |
| $K_{ijk}$ | Half-saturation constant | mmol ml<sup>-1</sup> |
| $c_{ijk}$ | Conversion rate from metabolism to bacterial density | Cells mmol<sup>-1</sup> |
| $\mu_k$ | Mutation rate in species $k$ | Molecules cell<sup>-1</sup> day<sup>-1</sup> |
| $D$ | Dilution rate | ml day<sup>-1</sup> |
| $Q_i$ | Input concentration of substrate $i$ | mmol ml<sup>-1</sup> |
| $a$ | Cost of generalism parameter | $(E_{ijk})^a$ has units molecules cell<sup>-1</sup> |

The resource use of each species is defined by a matrix *E*_*k*_ that specifies the amount of enzyme per cell for each reaction converting substrate *i* into substrate *j*. Species can be specialists on one reaction (i.e. only one *E_ijk_*>0) or generalists on several. Constraints are used to define pathways for the whole system (fig. 1); for example, here unidirectional catabolism breaking input resources into derived smaller molecules is assumed (only *E*_*ijk*_ above the diagonal are positive and input resources have the lowest indices *i*). Pathways can branch (*E*_*i,,k*_ is positive for multiple *j*) or coalesce (*E*_*,jk*_ is positive for multiple *i*). The total amount of enzyme produced per cell is assumed fixed across all species [31]: resource use varies because species allocate different proportions of their total metabolic enzyme to different reactions in a linear trade-off [32]. The effects of varying the trade-off between specialism and generalism is considered below. For simplicity, enzyme expression is assumed to be constitutive, i.e. there is no plasticity. Alternative modes of expression such as regulated resource switching could be investigated in future. Kinetics can vary across species and reactions: species with higher *v* for a given substrate grow more rapidly at high substrate concentrations, those with lower *K* more efficiently at low substrate concentrations.

The dynamics are then modelled by the following ordinary differential equations (ODEs), which sum the metabolism of each substrate in each species:

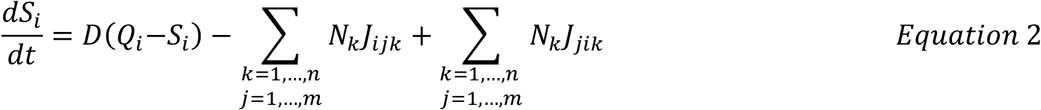

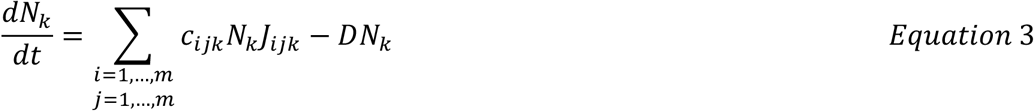

The terms for substrate concentrations represent dilution, metabolism of substrate *i* and production of substrate *i* in turn. The terms for population densities represent growth and dilution in turn. Parameter *c*_*ijk*_ is a constant converting metabolism of a given reaction into bacterial cells, which encapsulates both ATP produced per molecule of substrate and the proportion of ATP devoted to growth [28]. Growth on multiple substrates is assumed to be additive [30, 33] – substrates are alternative resources that can be metabolised without interference (empirically reasonable, especially in low nutrient systems, [33, 34]. There are *n*+*m* state variables and up to *2nm(m-1)* +*m*+*1* parameters in the fully specified catabolic model (only parameters above the diagonal for each matrix *v, K, E* and *c*). Parameter complexity is simplified by constraining pathways (i.e. assigning many values to 0) or by restricting which parameters are free to vary among species and reactions.

#### Analytical solutions

With continuous flow and no evolution, species growth and metabolism leads to steady-state concentrations and species densities [30, 35]. Steady-state solutions for a linear pathway performed by a set of species each specialized to metabolise just one substrate can be calculated for the Monod model (Appendix, fig. 2). For substrate *i* used by species *i*, steady-state values of the state variables are:

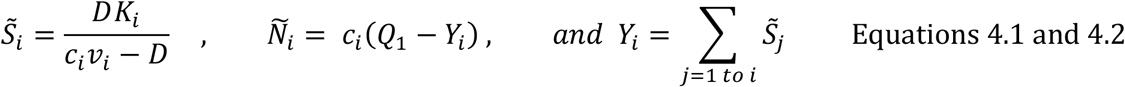

for all substrates except the final substrate (Appendix I, [29]). These solutions are stable as long as species *i* grows faster than it is washed out by the outflow (fig. 2D). If so, the final substrate, *m*, which is not metabolized, has steady-state concentration 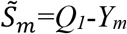. These expressions can be used to calculate various measures of community functioning and to design possible interventions to enhance functioning (Table 2, Appendix, fig. S1). Here, I focus on the steady-state concentrations of input, intermediate and final substrates.

**Table 2.** Example metrics of functioning that can be calculated for a chemostat model with a linear pathway performed by a set of specialist species (see Appendix for derivations).

| Scenario | Metric of functioning | Possible interventions to enhance functioning |
| --- | --- | --- |
| Bioremediation | Reduce concentration of toxic input substrate in the outflow,<br>$\tilde{S}_1 = \frac{DK_1}{c_1 v_1 - D}$ | Introduce a species with higher $c_1 v_1$ or lower $K_1$, which is feasible as it would outcompete incumbent species [36].<br>An easier solution is to reduce $D$. |
| Waste-water treatment | Decompose the greatest amount of input substrate per unit time,<br>$D(Q_1 - \tilde{S}_1)$ | Set dilution rate to intermediate optimum value for production, given by<br>$D_{opt} = \frac{c_1 v_1 (K_1 + Q_1 - \sqrt{K_1(K_1 + Q_1)})}{K_1 + Q_1}$ |
| Animal gut | Increase concentration of intermediate substrate (e.g. short chain fatty acids),<br>$\tilde{S}_i = \frac{DK_i}{c_i v_i - D}$ | Introducing a species with lower $c_i v_i$ or higher $K_i$ is unfeasible, as it would be outcompeted by incumbent species and could not invade. Instead could increase mortality of species $i$, e.g. with selective antibiotics or phage.<br>Easier option is to tune $D$ to be slow enough to allow earlier steps in pathway to be sustained but fast enough that product $i$ cannot sustain growth of a bacterial population, defined by:<br>$D < \frac{c_j v_j (Q_1 - Y_j)}{(K_j + Q_1 - Y_j)} \text{ for all } j < i \text{ but } D > \frac{c_i v_i (Q_1 - Y_i)}{(K_i + Q_1 - Y_i)}.$ |
| Bioreactor | Increase production of end product,<br>$\tilde{S}_m = Q_1 - Y_m$ | Increase concentration of input resource, or reduce $K_i$ or increase $c_i v_i$ of species performing any intermediate reaction |
|  |  | (the metabolic step to target with the biggest influence on final yield be calculated, see Appendix).<br>If feasible, shorten the pathway, e.g. by finding a species that converts substrate $i$ to substrate $i+2$ directly (under certain conditions, see Appendix).<br>Tune $D$ to the optimal rate for production. |

**Figure 2.**
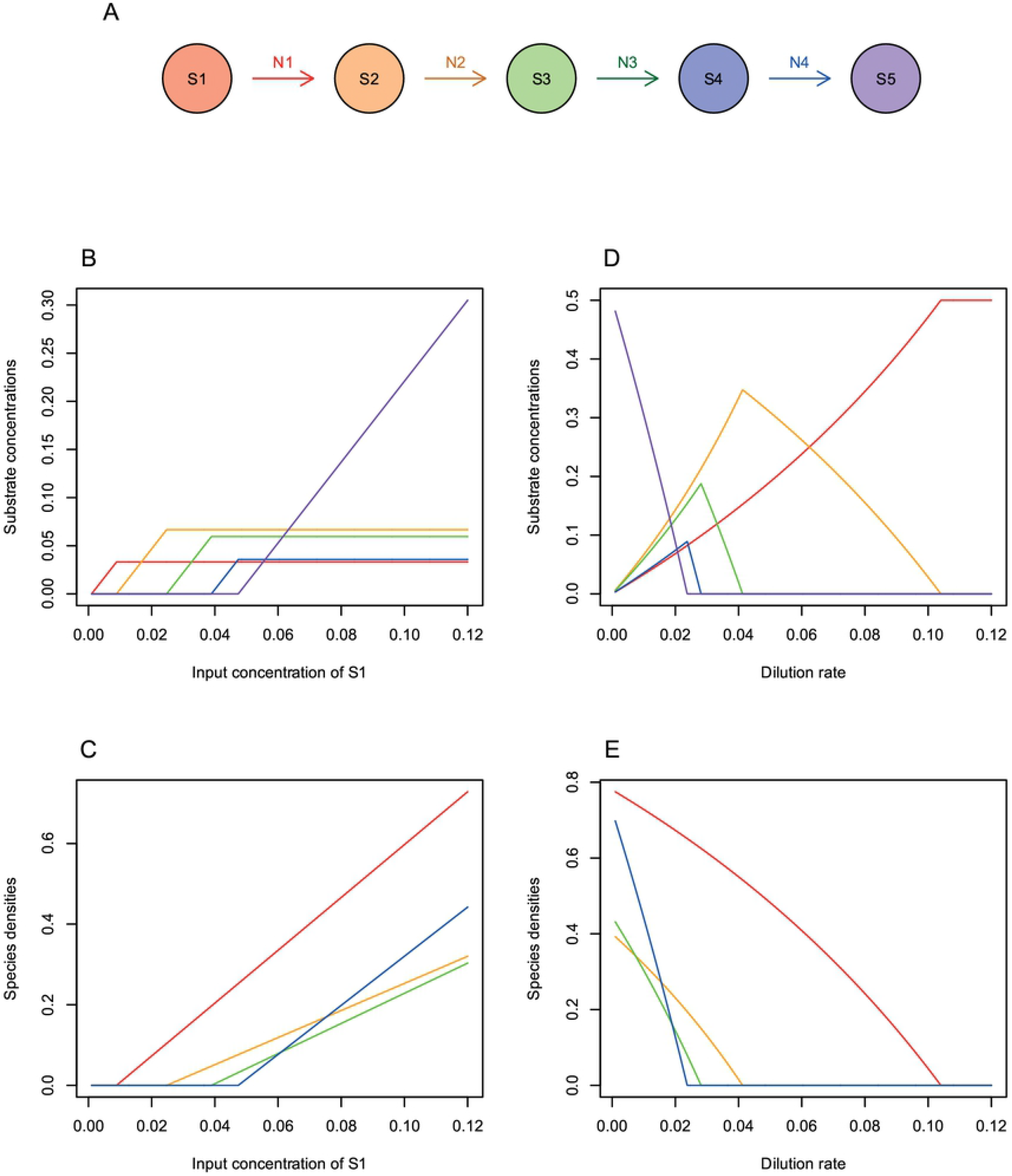
Analytical solutions for a linear pathway with specialist species and no evolution (A). Parameter values were: dilution rate, D=0.01 (in B and C); input substrate concentration, Q_1_ = 0.5 (in D and E); all v = 0.2; all K = 1; values of c chosen from a normal distribution with mean 1 and sd 0.2, = 1.56, 0.80, 0.89, 1.45 for each reaction in turn. Increasing the concentration of input resource increases allows successive intermediate substrates in the pathway to accumulate up to a maximum steady-state value (B), as species performing those steps are able to grow faster than the rate of washout (C). Reactions with higher biomass yields per molecule of substrate metabolized (e.g. red and blue) have lower steady-state concentrations of substrate and a faster increase in species density as input resources increase. Increasing the dilution rate has a reverse kind of effect: only earlier steps in the pathway can be sustained as dilution rates increase (D and E). For each intermediate substrate, there is a single dilution rate that maximizes its concentration at steady-state (D), which is the threshold below which the species that can use that substrate is able to persist (E).

### Evolutionary model

I added evolution to the chemostat model by assuming that species evolve through changes in their enzyme allocation (matrix **E**_**k**_ for species *k*) over time, modeled as the product of the genetic variance-covariance matrix (**G**_**k**_) and the selection gradient, which relates the mean growth rate of each species 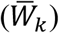 to changes in the expression level of each enzyme in turn:

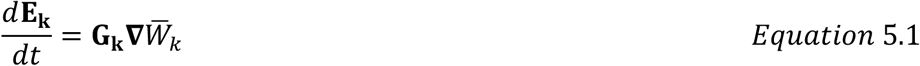

where the elements of

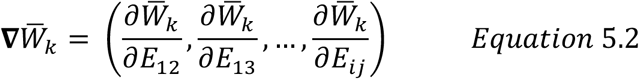

and 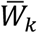 is the mean growth rate per cell per unit time [37], i.e. equation 3 divided by *N*_*k*_. The diagonal elements of matrix **G**_**k**_ are the heritable variation in enzyme expression, *μ*_*k*_. Under the constraint that total enzyme production is fixed and there are no other genetic correlations among enzymes, the non-diagonal elements are −*μ*_*k*_ /(*m*-1). For example, with 2 enzymes the genetic correlation between them is −1.0, i.e. as one increases in frequency the other decreases by the equivalent amount. Boundary constraints were included in the simulations so that changes to enzyme levels above one or below zero were truncated.

Equation 5 derives from quantitative genetics models of [38] as applied to clonal populations by [39]. It assumes large populations and a lack of substructure within species (e.g. diversification) such that changes can be modelled as changes in the mean phenotype of each population. It is chosen as the simplest model to allow changes in mean resource use without introducing additional structure in an already complex model (Appendix).

### Implementation

The model was simulated in the R statistical programming language using the lsoda function of the deSolve package. Code is available from github. Simulations were run for the ecological model (no evolution) and the evolutionary models in turn: initially for linear pathways with 1, 2 and 3 species, before implementing a branching and coalescing pathway performed by 8 species. Steady-state solutions for models with 1 to 3 species for the linear growth function were explored using SageMathCloud and Collaborative Calculation in the Cloud (CoCalc). The effects of the packaging of enzymes among species were investigated by comparing models with generalists that express multiple enzymes to models with the same enzymes and values of kinetic parameters present, but with each enzyme restricted to a different specialist species.

## RESULTS

### Does packaging of enzymes into species matter for functioning? One non-evolving generalist

I first consider a single non-evolving generalist that metabolises two input resources with chemostat concentration *S*_*1*_ and *S*_*2*_. I assume that kinetic parameters of each enzyme are the same as in the specialists but the generalist allocates a fraction *E*_*1*_ of its total enzyme to substrate 1 and *E*_*2*_ to substrate 2. Outcomes are more complicated than with two specialists because dynamics of substrate 1 now also depend on the amount of growth on substrate 2 and vice versa. If *E*_*1*_ = *E*_*2*_ = 0.5, the input concentrations of each substrate are equal, and both enzymes have the same kinetic parameters, then 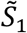 and 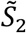 are the same for the generalist as for the two specialists (Fig. 3A). In all other cases, solutions differ between the generalist and specialist cases. For example, if *E*_*1*_ > *E*_*2*_, then 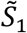 is reduced marginally compared to the specialist solution (for 0.5 < *E*_*1*_ < 1.0, fig. 3B), whereas 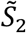 increases, reaching the input concentration *Q*_*2*_ when no enzyme is allocated to substrate 2 (fig. 3B).

**Figure 3.**
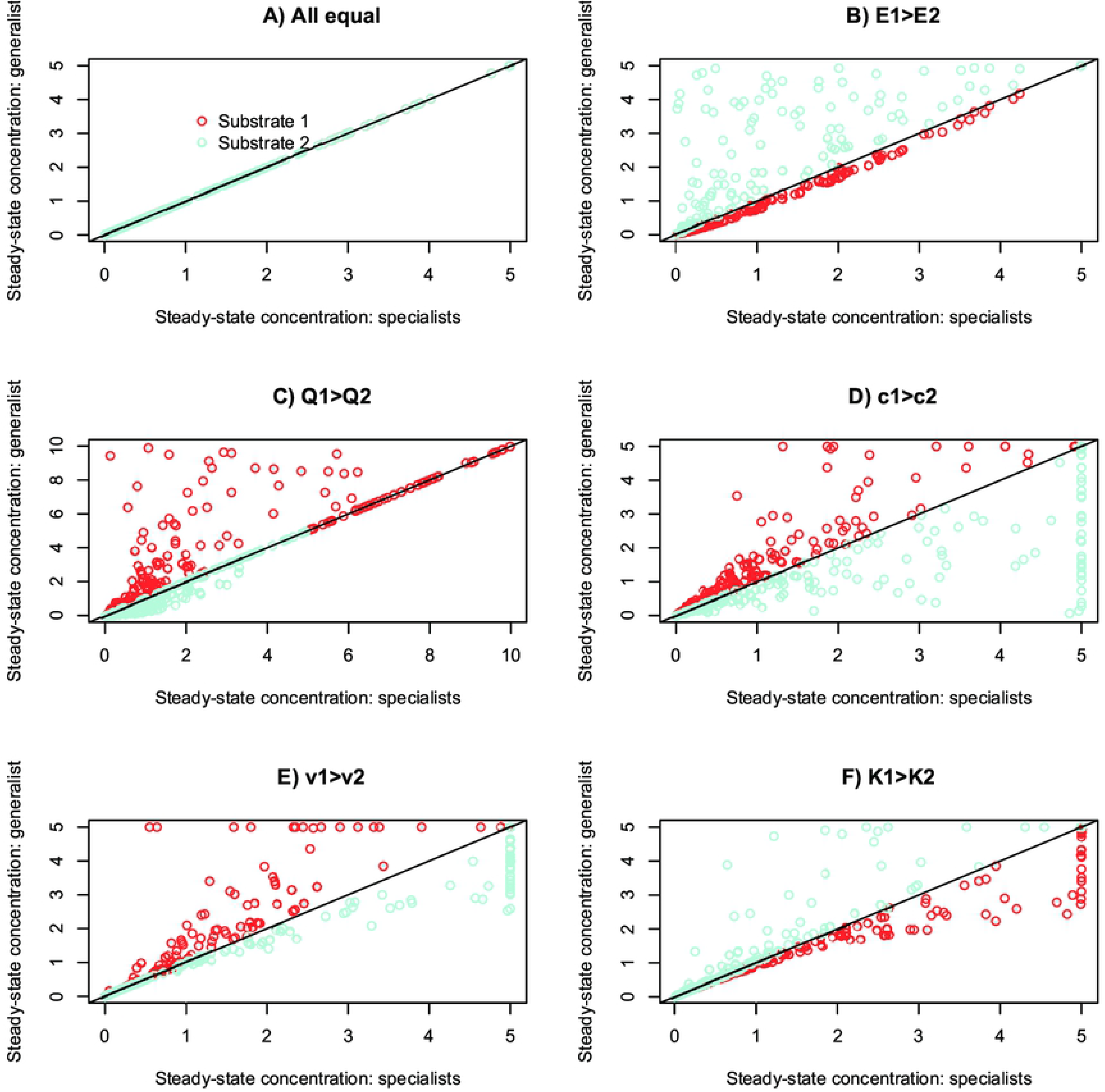
Comparison of steady-state solutions with 2 input resources both metabolized into a single waste product by either two specialists (X-axis) or one generalist species (Y-axes). Each point represents a separate run with parameter values chosen from uniform distributions (c=0.1 to 1, v=0.1 to 1, K=0.01 to 5, D=0.01 to 0.05, Q= random partition, i.e. broken stick, of 10 between the two substrates, E = random partition of 1.0). Parameters were fixed to be the same for substrate 1 and 2 except for the parameter varied in each panel as described in titles. The same parameters were used for matching specialist and generalist runs represented by a single point in order to compare the effects of enzyme packing into species. Simulations ran for 2000 time units.

Other parameters have consistent effects: the more profitable substrate for growth (either higher *c* or *k*, or lower *m*) is metabolised less by the generalist than by two specialists, whereas the less profitable substrate is metabolised more (Fig. 3D,E,F). This is because growth generated by metabolism of the more abundant resource increases total enzyme for the rarer resource, *NE*_*2*_, above levels that could be sustained by a specialist; but devoting half the cellular enzyme to the less abundant resource reduces metabolism of the more abundant substrate. In some cases, one of the specialists cannot persist but a generalist can (vertical sequence of blue points at 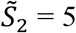, fig. 3E), and in others specialists can persist but a generalist cannot (horizontal sequence of points at 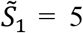 fig. 3E). Clearly the packaging of enzymes into different species alters functioning in this simplest case. Similar results are obtained for a pathway in which substrate is metabolized into derived substrate 2, which itself can be metabolized for growth (Appendix, fig. S2).

### Does the packaging of enzymes into species matter? Two non-evolving generalists

The situation changes when 2 species coexist on 2 substrates. Now, irrespective of whether there are 2 specialists, 2 generalists, or 1 specialist and 1 generalist, the steady-state concentration of both substrates is the same as the specialist case,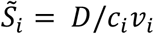, whenever both species persist. The only change in metabolic functioning occurs if one or both species cannot persist, in which case functioning is given by single-species solutions. Coexistence is determined by the following threshold enzyme allocation for the 2-input model with linear growth function (fig. 4):

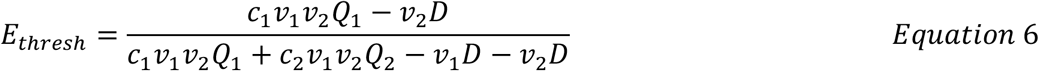

**Figure 4.**
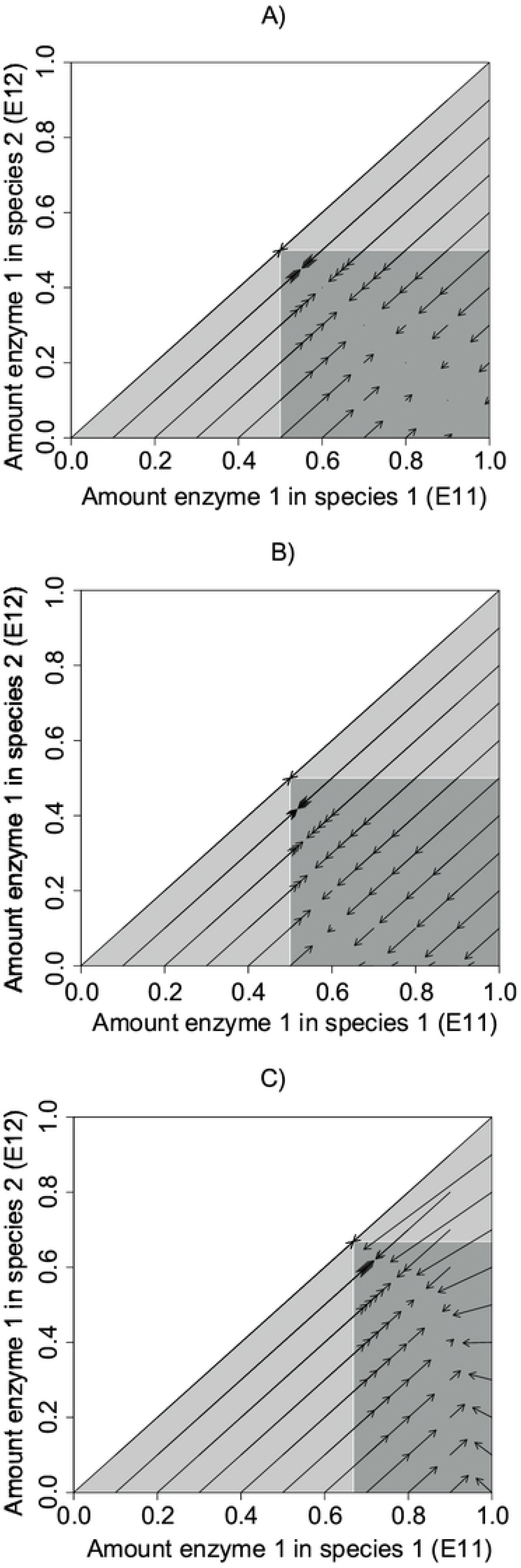
Evolutionary trajectories with 2 species and 2 input resources starting from different enzyme allocations. Species 1 allocates enzyme to substrate 1 more than species 2 does (E11>=E12). (A) All parameters equal, v=0.5, c=0.5, Q_1_=Q_2_=5, D=0.02. (B) With different starting densities, N1=0.5, N2=0.1. (C) With different conversion parameters, c_1_=1.0, c_2_=0.5. In each case, species evolve towards a region of neutral equilibrium defined by the threshold in equation 6, but the final enzyme allocation and densities depend on starting values. The dark grey region also indicates the region of coexistence of 2 species for an ecological model with no evolution. The lower light grey triangle indicates the region where species 1 alone persists and the upper light grey triangle indicates the region where species 2 alone persists.

If one species devotes more enzyme to substrate 1 than this threshold (e.g. *E*_*11*_ > *E*_*thresh*_) and the other species devotes less enzyme to substrate 1 than the threshold (i.e. *E*_*12*_ < *E*_*thresh*_), then the species can coexist. When all enzyme parameters and input concentrations are the same for both substrates, *E*_*thresh*_ is 0.5: one species needs to be more specialized on substrate 1 and the other more specialized on substrate 2, but even a small divergence from 50:50 allocation is sufficient for coexistence. Otherwise, if both species devote more enzyme to substrate 1 than this threshold (*E*_*11*_ > *E*_*thresh*_ and *E*_*12*_ > *E*_*thresh*_) then the species with the lowest allocation to substrate 1, i.e. the more generalist, will persist alone, and functioning collapses back to the single generalist case described in the previous section (fig S3). A similar but reverse outcome occurs if both species devote more enzyme to substrate 2 (*E*_*11*_ < *E*_*thresh*_ and *E*_*12*_ < *E*_*thresh*_, fig S3). The conclusions also apply to a pathway with 1 input substrate and 1 derived substrate, to 3-species coexistence (Appendix), and to the model with a Monod growth function (fig S4). Comparing 2 specialists with 2 generalists, functioning is therefore independent of how enzymes are packaged among species as long as both species are able to persist in both cases.

### The effects of evolution in one and two species models

Steady-state solutions of the linear growth model can be obtained for a single generalist evolving in its use of 2 input substrates according to equation 4.

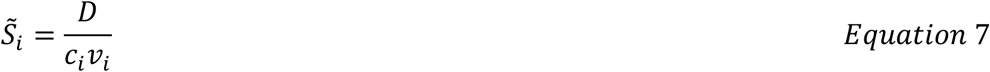

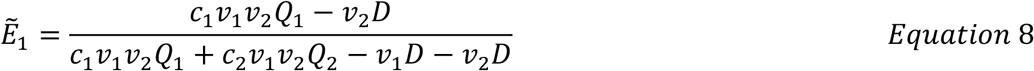

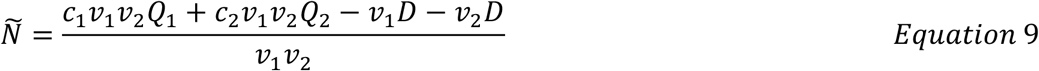

Evolution therefore simplifies the solution compared to the non-evolving model and yields a steady-state enzyme allocation with substrate solutions equivalent to 2 specialist species. The species devotes more enzyme to the resource with the highest balance of growth yield and rate (i.e. higher value of *c*_*i*_*v*_*i*_*Q*_*i*_ − *D*). A specialist on substrate 1 evolves when growth on substrate is unsustainable (i.e. 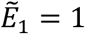 when *Q*_2_ ≤ *D*/*c*_2_*v*_2_ as long as growth on substrate 1 is sustainable, i.e. *Q*_1_ > *D*/*c*_1_*v*_1_). A specialist on substrate 2 evolves with the reverse conditions. A similarly tractable result is obtained for a single generalist adapting to 1 input substrate 1 and 1 derived substrate: growth is sustained as long as growth on the input substrate is viable; enzyme is always allocated to the derived substrate 2 as well for all viable solutions; and there is an inherent benefit towards specializing on the input resource, even if the growth yield on both substrates is equivalent (Appendix, see also [40]). [41] present general results for optimal allocation of enzyme in a single species using metabolic control theory.

With two evolving species present, there is a region of neutral equilibrium meeting the criteria for coexistence of two species outlined above: one species has *E*_*11*_ > *E*_*thresh*_ and *E*_*12*_ < *E*_*thresh*_, the other the opposite. Systems evolve towards enzyme allocations within the equilibrium region determined by starting enzyme allocations and densities (fig. 4), and both species persist as long as both substrates deliver sustainable growth. The outcome can either be partly complementary generalists or one specialist and one generalist. Once at steady-state, all systems deliver the same metabolic functioning as two specialists or a single optimal generalist, irrespective of starting conditions. There is an eco-evolutionary interaction since the evolution of species 1 depends on the starting density and enzyme allocation of species 2 and vice versa. Once a solution with *S*_1_ = *D*/*c*_1_*v*_1_ and *S*_2_ = *D*/*c*_2_*v*_2_ is reached, the growth rate of a perfect generalist species or two specialists invading at low frequency is 0, showing that the solution is evolutionarily neutral (Appendix). Similar dynamics are observed for the 1 input, 1 derived substrate model and for the model with Monod growth (fig S4).

Evolution therefore facilitates coexistence of 2 generalists and reduces competitive extinction. As a result, there is considerable functional redundancy since, within the bounds outlined above, sets of species with different enzyme allocations can provide the same metabolic functioning.

### Ecology, evolution and functioning in a decomposition pathway: 8 species, 11 substrates

To illustrate metabolic functioning of a more diverse community, I simulated communities with 8 species performing a decomposition pathway inspired by fermentation of indigestible polysaccharides in the human colon (Fig. 5, [42]). Two input substrates (starch and inulin) are degraded via linear pathways to a set of shared intermediates (lactate and the short-chain fatty acid acetate) and 3 terminal metabolites (the short-chain fatty acids butyrate and propionate plus waste gas, representing CO_2_ and methane, which I assume is not metabolized). Starch enters at higher concentration than inulin (3 versus 2 units of concentration). The pathway could also represent breakdown of two input resources (e.g. cellulose and chitin) by a soil or aquatic bacteria community [43]. My aim is to investigate in theory how species packaging and evolution affects functioning, rather than simulate a real pathway, which in the gut contains many more steps [42].

**Figure 5.**
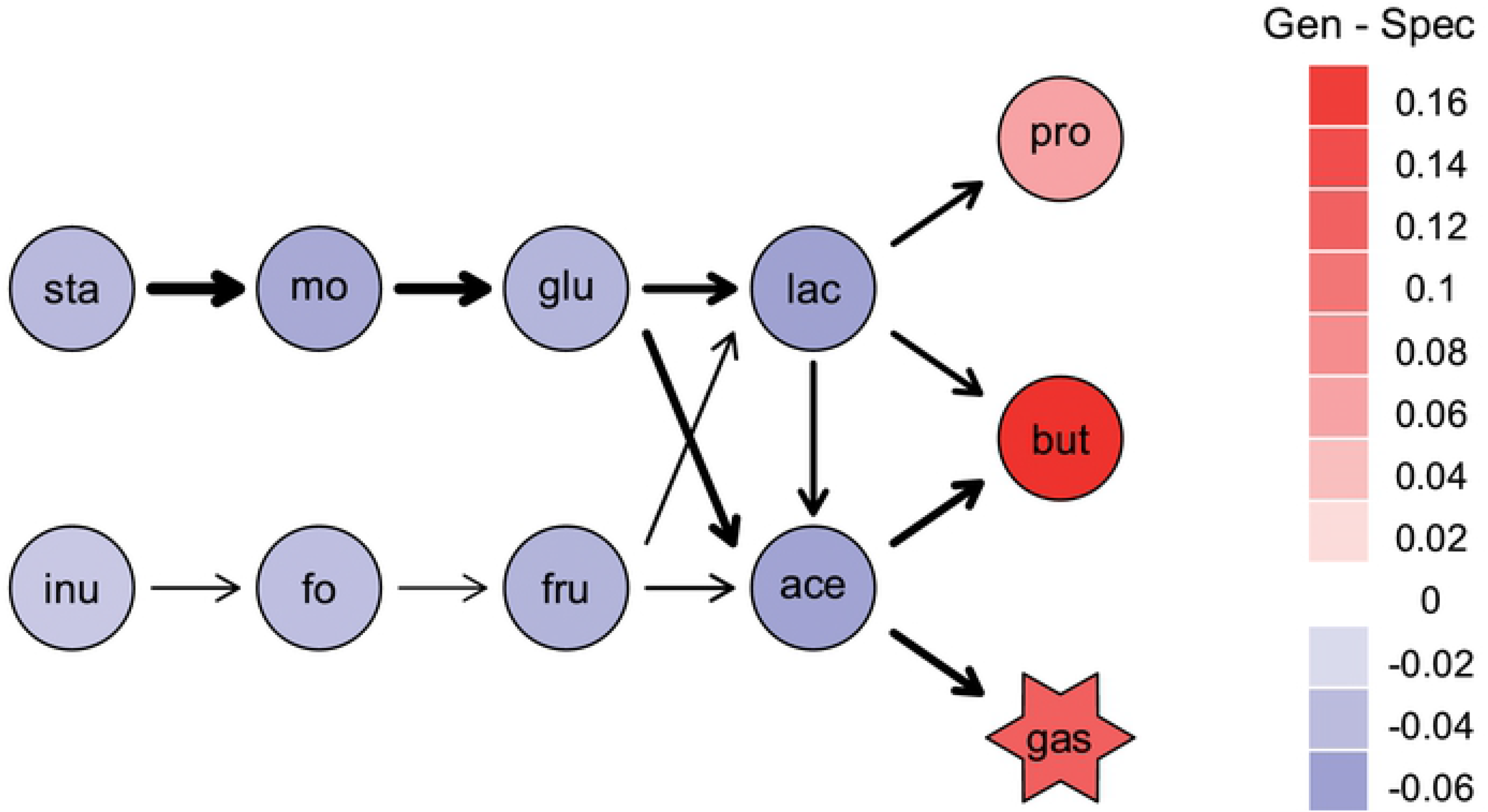
A decomposition pathway inspired by fermentation of indigestible polysaccharides in the human gut by the microbiome. Simulations of the ecological model with 8 species were run in pairs: one version with a set of specialists degrading each input and intermediate metabolite in turn and one version with generalists with random complementary allocation of enzymes for each metabolic step among species (so that each enzyme summed to 1.0 across the set of species, and enzyme allocation summed to 1 within each species). Parameter values were drawn from uniform distributions (between 0.13 to 1 for c, v and K parameters, and between 0.01 to 0.05 for D). Values for each enzyme were the same within a specialist versus generalist pair of simulations, but varied among 1000 replicated runs of those pairs. Red = higher steady-state concentrations of a metabolite for generalists than specialists, on average across 1000 replicates. Blue = lower concentrations for generalists than specialists. The thickness of arrows is proportional to the difference in average allocation of enzyme to that step between the starting species of generalists and the final surviving species: values ranged from 0.008 for starch to malto-oligosaccharides to −0.006 for fructo-oligosaccharide to fructose.

Again, I randomly allocated enzyme among species so that starting inoculum contained the same amount of each enzyme in every run. Enzyme parameters were chosen at random from uniform distributions set to allow general persistence of the diverse system (*c* = 0.13 to 1, *v* = 0.13 to 1, *K* =0.13 to 1, *D* = 0.01 to 0.05) and Monod growth was used since analytical solutions were unfeasible. I assumed that both steps out of a branching point in the network have the same rate parameters and yields, and starting species using that metabolite were allocated equal amounts of each enzyme (e.g. 33% of enzyme breaking down lactate is assigned to each of the three breakdown steps to produce propionate, butyrate and acetate in turn).

#### Ecological model

With the chosen range of parameter values, all 8 species survived in most runs of the ecological model with specialists (874/1000, average number of species surviving = 7.8 ± 0.8 sd). With generalists, fewer species survived: only rarely did all 8 species survive (40/1000, average number of species surviving = 5.3 ± 1.3 sd). The most abundant species comprised an average 42±12% of total bacterial yield in the generalist communities versus 24±6% of yield in the specialist communities. Surviving species on average allocated more enzyme to the starch input pathway than starting species, because of the higher influx concentration of starch than inulin, and to acetate degradation, because more energy flows through acetate than other intermediates in this pathway (arrow thickness, Fig 5). Survivors were also marginally more distinct from each other in enzyme allocations (mean pairwise Euclidean distance higher in 622/1000 cases, 2.8% higher on average) and slightly more specialist (mean allocation to leading enzyme higher in 573/1000 cases, 1.8% higher on average) than the starting species.

The steady-state of input and intermediate substrates was on average lower for the generalist community than for the specialist community (colour shading Fig. 5, fig. S5). In contrast, the concentration of terminal metabolites was on average higher for generalists as was total bacterial biomass (higher for generalist communities than specialist communities in 857/1000 cases, 5.8% higher on average, fig. S5). Together this indicates that the whole pathway ran at a faster rate with generalists than specialists: generalists sustained higher community-level concentrations of enzymes for less profitable steps than specialist communities could.

Transient dynamics also varied (fig. S6). On average, the generalist communities took 72% longer to reach steady-state for input substrates than specialist communities but were 20% quicker to produce terminal substrates. This occurred because growth on the input substrates increases concentrations of enzymes for later substrates in the generalist community, whereas specialists on later substrates only grow once those substrates are present in high abundance.

#### Evolutionary model

Allowing species to evolve changes the outcomes. Now 8 species survived in nearly all runs (998/1000) and metabolic functioning converged towards the corresponding specialist run (fig. 6, fig. S7). Exceptions arose when not all specialist species could survive, i.e. because one step did not yield sufficient energy to outgrow dilution rate. The evolving generalists retained some enzyme allocation to those substrates because they do provide energy for growth, just not enough to sustain a specialist. In many of those cases, all 8 species evolved to have near identical phenotypes, i.e. the optimum allocation for a single species utilizing this pathway (Fig. S8). Even when 8 functionally divergent species survived (905/1000 runs) species tended to converge in enzyme allocation patterns (mean pairwise Euclidean distance 0.29±0.06sd versus 0.41±0.04sd for surviving generalists in the ecological model).

**Figure 6.**
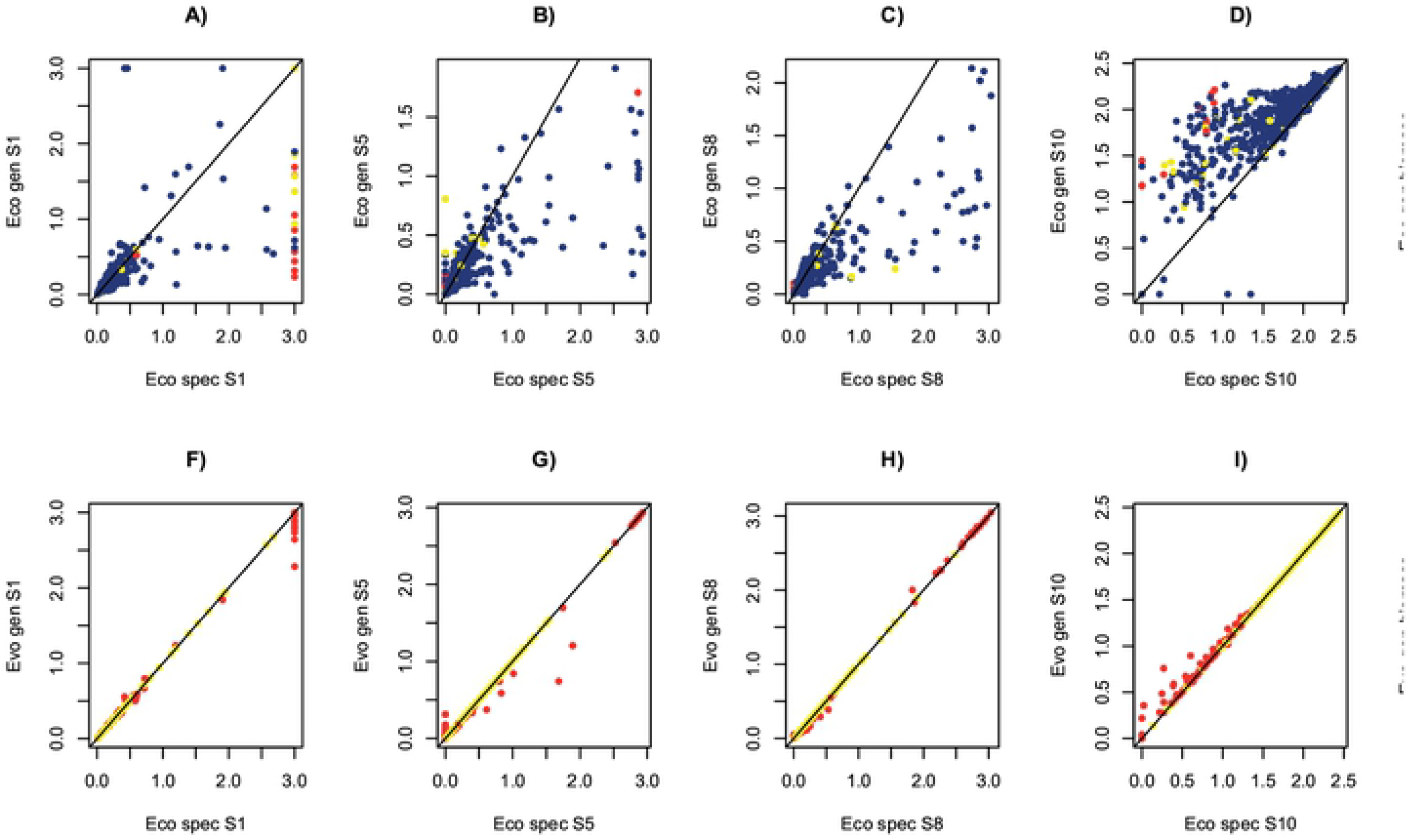
Simulations of the simplified gut pathway starting with 8 species. In each plot, the value for the generalist community is the Y axis and the value for the matching specialist community is the X axis. The top row (A to E) shows results for the ecological model and the bottom row (F to J) shows results for the evolutionary model. An example of an input substrate (S1, starch, A, F), intermediate substrate on input pathway (S5, glucose, B, G), intermediate substrate on converging pathway (S8, acetate, C, H) and terminal substrate (S10, butyrate, D, I), and the total bacterial biomass (E, J) are shown. Blue = more species survive in specialist run; yellow = same number of species survive; red = more species survive in generalist run. Each point represents a separate run with parameters chosen from uniform distributions (0.13 to 1 for c, v and K parameters, 0.01 to 0.05 for D, E of each species = random partition of 1.0 for the generalist model), which were the same for each enzyme for the matched specialist and generalist versions of each run. Simulations ran for 20000 time units, with 1000 replicated runs.

#### Evolutionary outcomes depend on the trade-off between specialism and generalism

The above models assume a linear trade-off that a generalist with equal allocation to two enzymes grows half as fast on each substrate (when in excess) as a specialist. To investigate the effect of stronger trade-offs, I introduced a parameter *a*, which I term the cost of generalism. Now the rate of metabolism for each reaction becomes

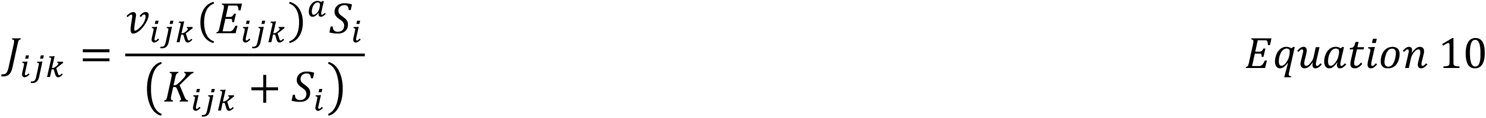

and the selection gradient for enzyme *ijk* becomes 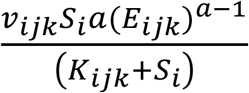. In a generalist, all *E*<1 and so *a*>1 causes a concave trade-off and the growth rate on each substrate is lower than with a linear trade-off. This reflects a cost of generalism, for example maintaining the machinery to express different enzymes, that increases as more enzymes are present.

A concave trade-off switches the balance towards specialism. Consequently, non-evolving generalist communities now display worse overall metabolic functioning than specialists, with higher concentrations of input and intermediate substrates and lower concentrations of terminal products (fig. 7, S9). Evolution now improves functioning relative to a non-evolving community of generalists (fig. S10), and species diverge in their enzyme profiles (fig. 8). Simulations started with specialists remained as strict specialists rather than converging as they did with a linear trade-off. The evolving community did not reach the level of functioning of a community of strict specialists, however, over the timescale of the simulations. There remain constraints therefore in reaching enzyme allocations across species that deliver optimum community-level functioning.

**Figure 7.**
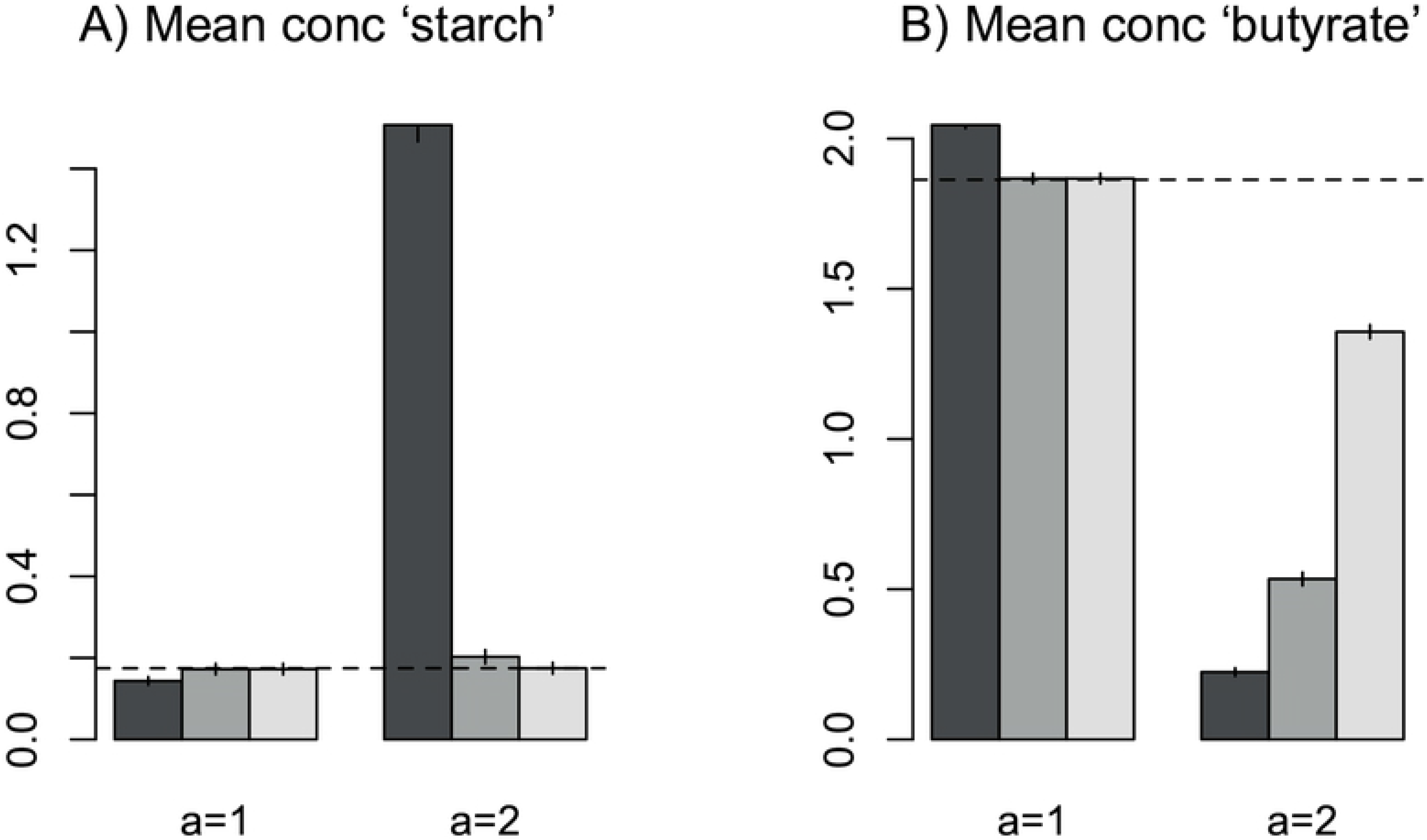
Average values of measures of functioning across trials with a linear trade-off for generalists (a=1) versus a concave trade-off (a=2). A) Mean concentration of substrate 1, i.e. input starch. Higher values = less efficient degradation. B) Mean concentration of substrate 10, i.e. butyrate, a beneficial terminal product. Values are shown for non-evolving generalists (dark grey), evolving generalists (mid grey) and evolving communities that started as specialists (light grey). Parameter values assigned as described in figure 6. Standard errors across 1000 replicate runs are shown.

**Figure 8.**
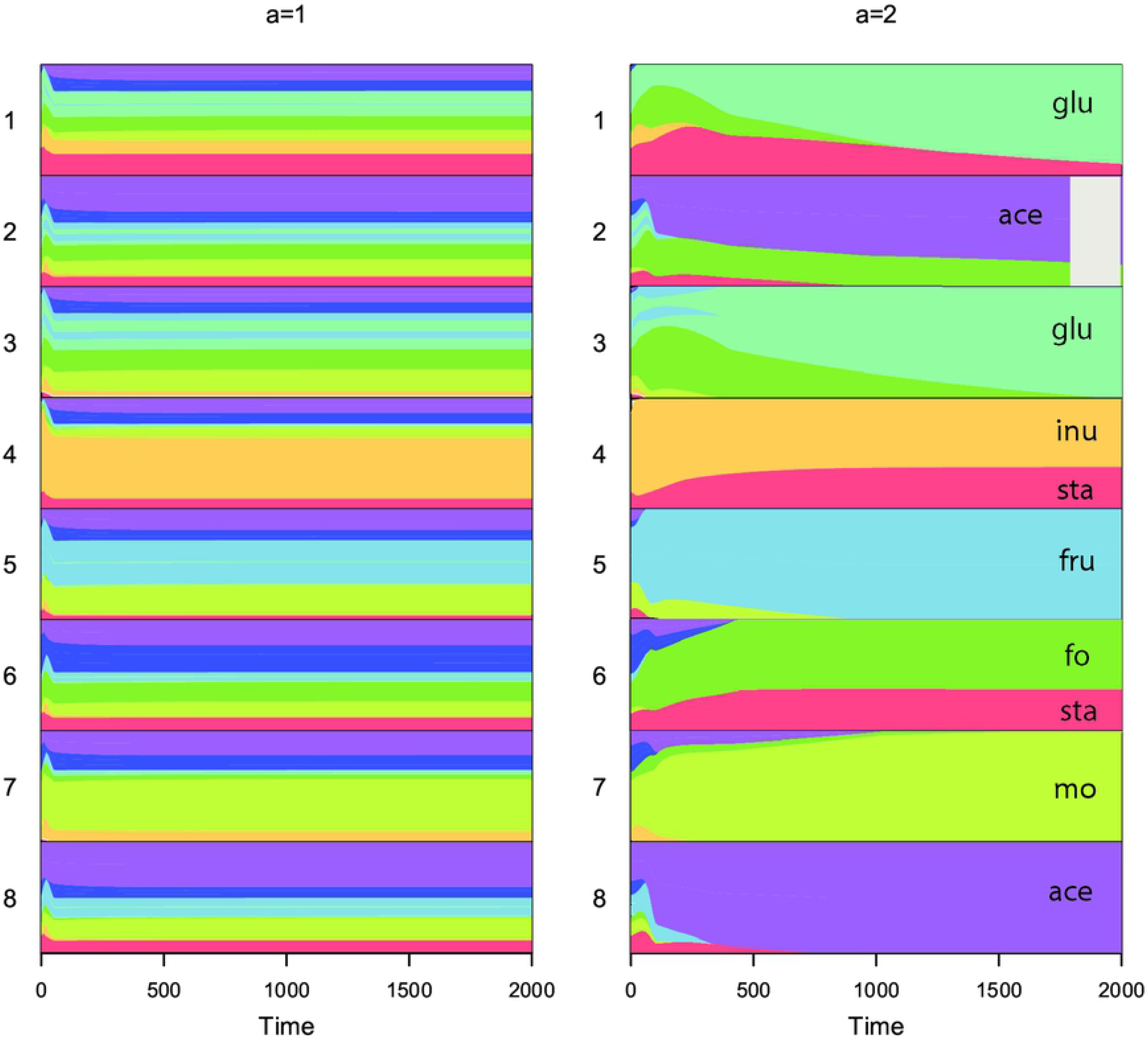
Evolution of enzyme allocations across the 8 species for one exemplar run over the first 2000 time steps with a linear trade-off for generalists (a=1) or a concave tradeoff (a=2). The run started in both cases with a generalist community. Each colour represents the proportion of enzyme allocated to metabolizing a different substrate (labels under a=2 correspond to labels of substrates on pathway in figure 5). With a concave trade-off, species evolve to be more specialists (more of one colour) and partition resources (species tend to have different colours). Species 2 went extinct within the first 2000 time steps with a=2. Parameter values were assigned as described in figure 6, just one exemplar run shown.

## DISCUSSION

The packaging of traits into species was important for overall functioning. A community of generalists functions differently from a community of specialists, because packaging introduces constraints on outcomes. Generalist communities can sustain metabolic steps that are not profitable enough for a specialist to grow faster than dilution, but in doing so generalist species lose out from additional growth on more profitable substrates. Transient dynamics also depend on enzyme packaging: generalist communities are slower to respond to an initial excess of input substrate, but quicker to perform linear decomposition pathways because enzyme levels for later steps are ‘primed’ by earlier ones.

Nonetheless, the models also predict considerable functional redundancy. Functioning converged on the same outcome whenever enzyme concentrations were able to reach optimum system-level concentrations: either by ecological changes in species densities or evolution of enzyme allocations. A range of species densities and evolutionary endpoints yielded the same metabolic functioning: there were multiple ways of packaging enzymes to yield identical functioning. There are reasons to suspect, however, that real communities are less redundant than found here. First, functional redundancy was only extensive when there was a weak cost to generalism (i.e. a linear trade-off). Second, I assumed that an enzyme’s kinetic parameters are the same irrespective of which species it is found in. If some species have faster reactions or lower saturation constants than others, this would shift competition in their favour. Third, I assumed that all species can evolve all patterns of enzyme allocation, which is unlikely. It is therefore likely that species matter even more in real communities than found here.

Evolution generally resulted in a less productive community that yielded lower levels of end products than a non-evolving community of generalists, when a linear trade-off for generalism was assumed. With a concave trade-off, evolution now improved metabolic functioning relative to the non-evolving generalist case, but it still did not always lead to perfect specialism, which yielded the best overall metabolic pathway rate in these conditions. These results generate predictions of when evolution should improve or reduce functioning of a set of species encountering a new environment, which depends on starting phenotypes as well as the selection gradients on changing enzyme profiles. They also confirm that optimum functioning of the whole community need not be selected for within species [26]. Models based on assuming optimality of the whole system might not be applicable to species communities: instead, models are needed that explicitly consider dynamics and progress towards final states.

The model incorporates eco-evolutionary interactions [44–46]. Species evolution depended on the traits of other species, and species densities were affected by evolution. I only modelled one mutation rate here, but because starting conditions affected final outcomes, the speed of evolution would change the outcome (e.g. lengthening or shortening trajectories in fig. 4). In the model with a linear trade-off for generalists, species tended to converge in trait values. Again, this likely reflects the frugal approach to parameters and broad equivalence of species except enzyme allocation. All species experienced the same selection pressure at each time based on resource availability, but outcomes differed because of different starting phenotypes. If species differed in enzyme kinetic parameters as well, then selection pressures would vary across species and lead to a wider range of outcomes.

How do these predictions compare to experimental evidence? Gravel et al. [47] evolved a set of species to be either more specialist or more generalist in turn and then constructed communities from them. The generalist communities displayed greater productivity than specialist communities (measured over 72 hours, which was too short for further evolution, hence equivalent to non-evolving simulations), which matches predictions here for the linear trade-off for generalism.

There is also some evidence for species evolving towards convergent resource use. Foster and Bell [48] found that random pairs of wild species isolated from tree-hole communities tend to have negative interactions indicative of overlapping resource use. These findings presented a paradox as to how species coexist in nature. One answer is that many other environmental parameters vary spatially or temporally and species grow differentially with respect to those. The model presents an alternative: species coexist in a near neutral equilibrium shaped by evolutionary responses to resource competition. Four studies of coevolving species in laboratory microcosms, however, find results more consistent with a concave resource trade-off: species evolved resource partitioning over time leading to increased community productivity [27, 43, 49, 50].

Clearly more quantitative evaluation is required to investigate such cases. The models do not currently match experiments in detail. I assumed unusual starting conditions that complementary generalists are present, in order to investigate the effects of species packaging while holding other variables constant. This assumption is unrealistic for arbitrary starting species. Another difference is that I assumed constant resource supply whereas experiments used serial transfer or growth in batch, which causes fluctuating levels of input versus derived resources [30]. These modifications could be investigated in future.

The model provides guidance for predicting microbial functioning from genome data. Under most conditions, single cell genomics or long-read sequencing technology would be needed to infer which genes are found together within genomes; a metagenomics gene-level approach would not be sufficient to infer overall function. More challenging still, replicates show that functioning depends greatly on quantitative parameter values, not just enzyme presence or absence. Genome data do not readily yield estimates of kinetic parameters for enzymes.

In principle, the model could be parameterised for quantitative predictions. Species’ growth rates can be measured on media of each substrate to estimate kinetic and conversion parameters. RNA sequencing could reveal which enzymes are expressed during growth on complex media [51]. Selection experiments in monocultures could estimate rates of evolution and genetic correlations between use of alternative resources [52]. A parameterised model could then be simulated to see if it predicts responses in an experimental community, followed by exploration of extra processes needed to improve model fit. Alternatively, with whole communities, a combination of modelling, experimental perturbation [53], and time series of genomic, transcriptomic and metabolic data [54, 55] could be used to estimate key dynamic quantities. For example, does a non-evolving model parameterised at an early time-point adequately predict dynamics at a later time point or is evolution of species traits needed to explain later dynamics?

To conclude, although there are theoretical scenarios in which packaging of enzymes makes little difference to overall functioning (specifically with evolution and a linear trade-off), it is more likely that the packaging of enzymes among species does effect overall functioning. Species-based ecological and evolutionary models are vital in order to track bacterial dynamics and functioning and to design interventions for enhancing beneficial functions.

## ACKNOWLEDGMENTS

Thanks to Ana Gomez and Michael Schmutzer for providing some code used in simulations and Tom Bell, Glenn Gibson, Laura Johnson, Russ Lande, Sarah Otto, Dan Reuman, Richard Sheppard for discussion and comments. This work was supported by NERC grant NE/K006215/1, a Leverhulme Trust fellowship and a Visiting Professorship at the University of British Columbia in 2013.

## SUPPORTING INFORMATION CAPTIONS

Appendix - additional explanation and derivation of the model

**Figure S1**. For a linear pathway of 4 specialists degrading a single input resource: (A) the amount of substrate 1 degraded per unit time, i.e. 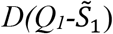 (B) The amount of each derived substrate produced per unit time, which is 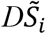 below the threshold for washout of species *i* and *D*(*Q*_*1*_− *Y*_*i*_) above the threshold. Parameter values were: input substrate concentration, *Q*_*1*_ = 0.5 (in D and E); all *v* = 0.2; all *K* = 1; values of *c* chosen from a normal distribution with mean 1 and sd 0.2, = 1.56, 0.80, 0.89, 1.45 for each reaction in turn. Note the maximum production rate of substrate 2 (orange) occurs at a higher dilution rate than the threshold for persistence of species 2 (which is at the inflection point on the orange curve), but for all other substrates the optimum dilution rate is at the threshold for persistence of the corresponding species.

**Figure S2**. Comparison of steady-state solutions with 1 input and 1 derived substrate metabolized by either two specialists (X-axis) or one generalist species (Y-axes). Each point represents a separate run with parameter values chosen from uniform distributions (*c*=0.1 to 1, *v*=0.1 to 1, *K*=0.01 to 5, *D*=0.01 to 2, *E* = random partition of 1.0). Input concentration of substrate 1 was held fixed at 5 units. Parameters were fixed to be the same for substrate 1 and 2 except for the parameter varied in each panel as described in titles. The same parameters were used for matching specialist and generalist runs represented by a single point in order to compare the effects of enzyme packing into species. Simulations ran for 2000 time units.

**Figure S3**. Simulations of a model with 2 input substrates and 2 species. Steady-state concentrations of substrate 1 (A) and 2 (B) for 2 specialists (x-axis) versus 2 generalists (y-axis). Red = just species 1 survives; blue = just species 2 survives; black = both species survive. Species 1 is defined as the species allocating most enzyme to substrate 1, i.e. E11>E12. Steady-state concentrations for substrates 1 (C) and 2 (D) when the generalist species are able to evolve during the simulation. Each point represents a separate run with parameter values chosen from uniform distributions (*c*=0.13 to 1, *v*=0.13 to 1, *D*=0.01 to 0.05, *E* = random partition of 1.0). Simulations ran for 20000 time units.

**Figure S4**. Simulations of a model with 1 input and 1 derived substrate and 2 species. Steady-state concentrations of substrate 1 (A) and 2 (B) for 2 specialists (x-axis) versus 2 generalists (y-axis). Red = just species 1 survives; blue = just species 2 survives; black = both species survive. Species 1 is defined as the species allocating most enzyme to substrate 1, i.e. E11>E12. Steady-state concentrations for substrates 1 (C) and 2 (D) when the generalist species are able to evolve during the simulation. Each point represents a separate run with parameter values chosen from uniform distributions (*c*=0.13 to 1, *v*=0.13 to 1, *D*=0.01 to 0.05, *E* = random partition of 1.0). Simulations ran for 20000 time units.

**Figure S5**. Simulations of the ecological model of the simplified gut pathway starting with 8 species. Steady-state concentrations of substrates and biomass for specialists (x-axis) versus non-evolving generalists (y-axis). Blue = more species survive in specialist run; yellow = same number of species survive; red = more species survive in generalist run. Each point represents a separate run with parameter values chosen from uniform distributions (*c*=0.13 to 1, *v*=0.13 to 1, *d*=0.01 to 0.05, E = random partition of 1.0). Simulations ran for 20000 time units.

**Figure S6**. An example of transient dynamics in substrate concentrations of a corresponding generalist (solid lines) and specialist (dashed lines) community for a linear pathway of 3 species. Similar outcomes were observed for the 8-species community but a 3-species systems is shown here for clearer visualization. Parameter values for this example: *c*_*1*_=0.20, *c*_*2*_=0.33, *c*_*3*_=0.23, *v*_*1*_=0.28, *v*_*2*_=0.57, *v*_*3*_=0.24, *D*=0.045, *Q*_*1*_=6, species 1 E_*11*_=0.04, E_*21*_=0.36, E_*31*_=0.60, species 2 E_*12*_=0.65, E_*22*_=0.01, E_*32*_=0.34, species 3 E_*13*_=0.32, E_*23*_=0.63, E_*33*_=0.05.

**Figure S7**. Simulations of the simplified gut pathway starting with 8 species. Steady-state concentrations of substrates and biomass for specialists (x-axis) versus evolving generalists (y-axis). Blue = more species survive in specialist run; yellow = same number of species survive; red = more species survive in generalist run. Each point represents a separate run with parameter values chosen from uniform distributions (*c*=0.13 to 1, *v*=0.13 to 1, *D*=0.01 to 0.05, E = random partition of 1.0). Simulations ran for 20000 time units.

**Figure S8**. The mean Euclidean distance for enzyme allocation between species at the start of each run and at the end of the run allowing species to evolve. Red = not all specialists survived in the specialist run. Yellow = same number of species survived in the specialist and evolving generalist runs. The 1:1 line is shown: in every run, the species are more similar in their enzyme profile by the end than at the start.

**Figure S9**. Simulations of the ecological model of the simplified gut pathway starting with 8 species and a cost of generalism exponent *a*=2, i.e. a concave trade-off for generalism. Steady-state concentrations of substrates and biomass for specialists (x-axis) versus non-evolving generalists (y-axis). Blue = more species survive in specialist run; yellow = same number of species survive; red = more species survive in generalist run. Each point represents a separate run with parameter values chosen from uniform distributions (*c*=0.13 to 1, *v*=0.13 to 1, *D*=0.01 to 0.05, E = random partition of 1.0). Simulations ran for 20000 time units.

**Figure S10**. Simulations of the evolutionary model of the simplified gut pathway starting with 8 species and a cost of generalism exponent a=2, i.e. a concave trade-off for generalism. Steady-state concentrations of substrates and biomass for specialists (x-axis) versus evolving generalists (y-axis). Blue = more species survive in specialist run; yellow = same number of species survive; red = more species survive in generalist run. Each point represents a separate run with parameter values chosen from uniform distributions (*c*=0.13 to 1, *v*=0.13 to 1, *D*=0.01 to 0.05, E = random partition of 1.0). Simulations ran for 20000 time units.

## Notes

Funding: This work was supported by NERC grant NE/K006215/1, a Leverhulme Trust fellowship and a Visiting Professorship at the University of British Columbia in 2013.

The author declares no conflict of interest

